# Studies on trans-sutural distraction osteogenesis-related genes based on transcriptome sequencing

**DOI:** 10.1101/2020.04.11.037085

**Authors:** Baicheng Wang, Hongyu Xue, Haizhou Tong, Peiyang Zhang, Mei Wang, Yidan Sun, Zhenmin Zhao

**Author notes:** WANG Baicheng, XUE Hongyu have equal contribution to the article. Corresponding author: Zhao Zhenmin, Tutor, Chief Physician, Research Director: Craniofacial plastic surgery and regenerative medicine.

## Abstract

Trans-sutural distraction osteogenesis (TSDO) is an important approach to improve mid-face hypoplasia. In recent years, many studies have been carried out on physical mechanisms of TSDO; however, it’s specific cytological and molecular mechanisms are still unclear. In this study, we performed transcriptome sequencing analysis in Sprague Dawley rats at 1 and 2 weeks after suture osteogenesis and compared RNA expression levels between experimental and control groups. At one week, enrichment pathways were mainly up-regulated in muscle- and bone-related pathways. By contrast, pathways of the immune system showed a state of inhibition and down-regulation, especially for B cells; the main immune pathways showed significant down-regulation. However, two weeks later, the experimental group showed positive up-regulation of the pathways related to DNA synthesis and replication, cell cycle, and chromosome replication. At the same time, the immune pathways that were down-regulated in the first week were up-regulated in the second week. In other words, the up-regulated muscle- and bone-related pathways show opposite trends. The expression of bone- and myogenesis-related transcriptome was up-regulated and the immune-related pathways were down-regulated in the experimental group at 1 week. At 2 weeks, the pathways related to bone- and muscle were down-regulated, while those related to cell cycle regulation and DNA replication were up-regulated. These results suggest that musculoskeletal-related molecules may play an important role during suture osteogenesis at 1 week, and immune regulation may be involved in this process; however, at 2 weeks, molecules related to cell proliferation and replication may be a major role.

## 1. INTRODUCTON

Central skeletal hypoplasia is one of the common cranio-maxillofacial deformities in pediatric practice, which mainly affects a child’s facial appearance, oral and maxillofacial system function, and the development of mental health[1]. The current treatments mainly include, traditional orthognathic surgery, osteotomy traction osteogenesis, and TSDO. The first two techniques require osteotomy when a patient’s bones are mature. Because of trauma and bleeding due to the surgery, it is not conducive to correct depression of the mid-face of teenagers[2]. Suspension traction osteogenesis directly applies traction to the natural bone suture of the human body without osteotomy, having the advantages of less surgical damage and simple technical operation. In addition, bone sutures of teenagers are growth areas during skeletal development and have a strong growth capacity. Therefore, surgical trauma and risks introduced by osteotomy can be avoided by the application of traction to bones. In addition, it can effectively restore teenagers’ or children’s mid-face bone functions; this technique provides new techniques and methods for early, minimally invasive correction of midfacial hypoplasia in teenagers[3]. The mechanism of suture osteogenesis through the technique of suturing is complicated. It is known that mechanical stimulus is transformed into biological signals to stimulate the histological process of cell activation, bone matrix formation, calcification, and bone reconstruction. Many transcription growth factors including, Bone morphogenetic protein (BMP), basic fibroblast growth factor(bFGF), Transforming growth factor-beta (TGF-β) and participate in the osteogenesis process of TSDO, which promote the proliferation and differentiation of mesenchymal cells to osteoblasts, leading to the reconstruction of new bone. However, the specific process of TSDO has not been elucidated[4, 5]. In recent years, some studies have speculated that during the suture traction, a large number of new bones are generated in a short term under the tractive force. Mesenchymal stem cells (MSCs) play an important role in this process, but specific cell types and pathways are not known[6, 7].

We propose a first automatic continuous suture traction osteogenesis in the world to correct skeletal dysplasia in the midface. Since 2005, more than 80 TSDO operations have been successfully performed, and all the patients have achieved satisfactory clinical outcomes [3, 8]. We have also established an TSDO rat model to investigate the mechanism of suture traction. Our study has used transcriptome sequencing technology to explore genetic changes in rats during suture traction.

## 2. MATERIALS AND METHODS

### 2.1 Animals

All experimental animals were 4-weeks-old Sprague Dawley (SD) male rats (100 g, obtained from the Animal Experiment Center of the Peking University Medical Laboratory). All the experiments were approved by the Ethical review committee (LA2019177) Rats were kept in 12-h light-dark conditions in a closed environment. The breeding temperature was maintained at 22 ° C ± 0.5 °C, and the humidity was 40-60%.

### 2.2 Study design

Twenty-four rats were randomly divided into an experimental group (n = 12) and a control group (n = 12). The experimental group were operated upon employing a surgical method described by Akiko Sasaki et al[9] and our previous experience of such surgeries. We cut the skin to expose the maxilla and temporal bones. We made two holes (4 holes in total) on the left and right maxillary and temporal bones, so that the distance between the two holes was 12 mm (Figure 2 A) and placed a traction device in them (Figure 2 E). The control group underwent the same operation, but without the traction device being placed. The operation was performed as expected and no animals died (Figure 1).

**FIGURE 1.**
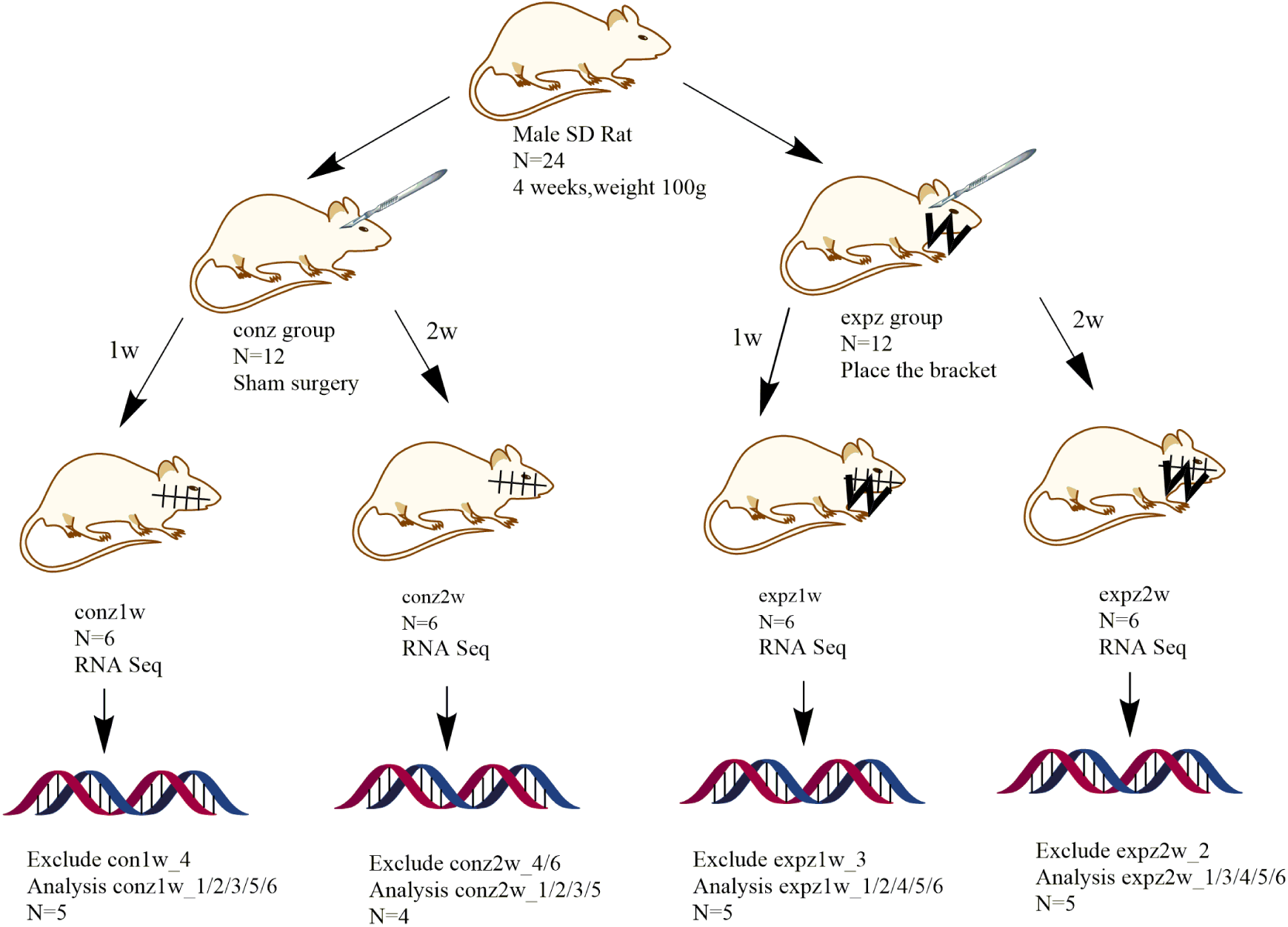
Experimental design. Sprague Dawley rats (n=12) were operated upon in the experimental group and in control group (n=12). Outliers (expz_3, expz2w_2, conz2w_4, con2w_6, and conz_4) were excluded.

**FIGURE 2.**
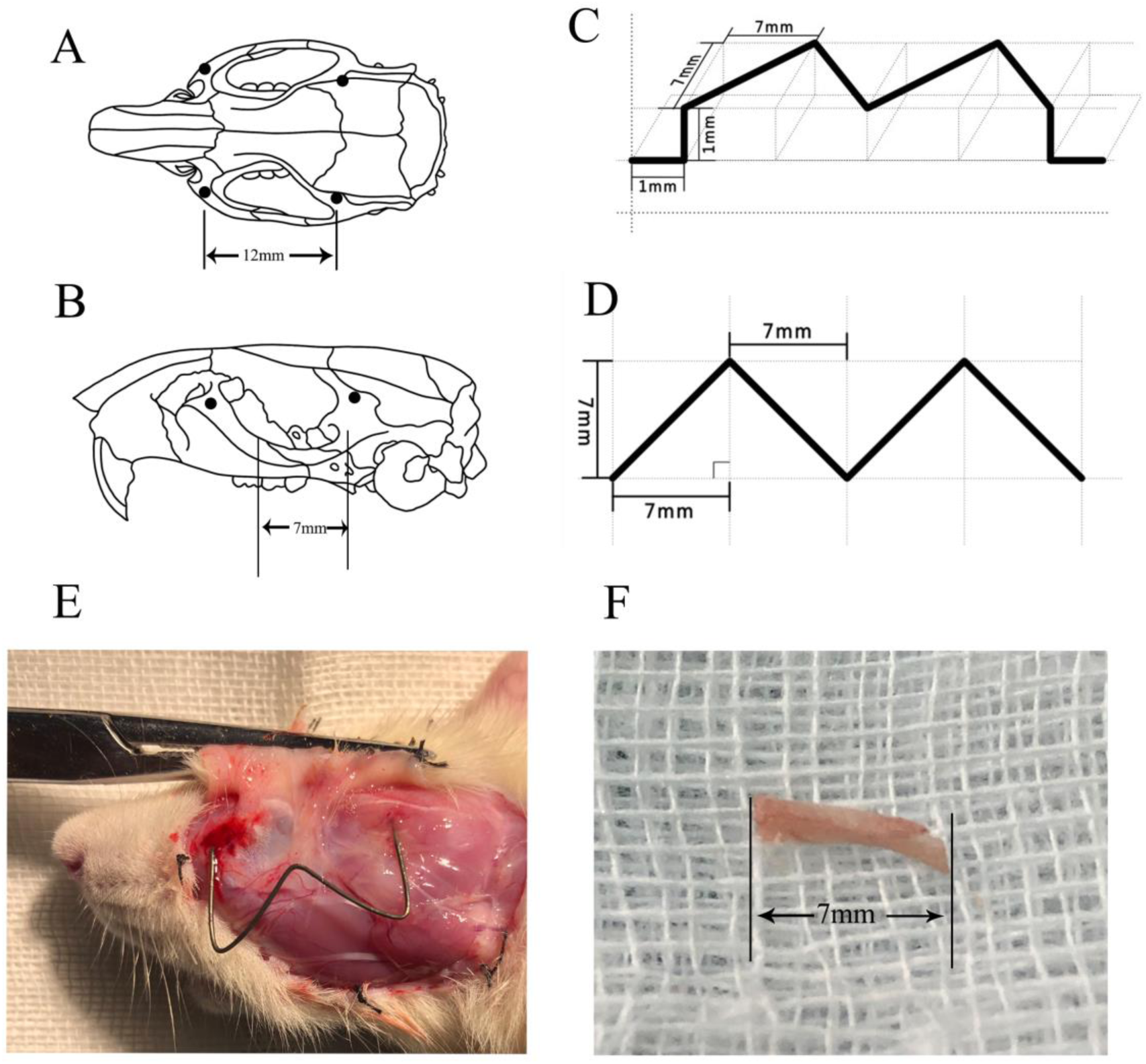
Plane and three-dimensional structure of the traction device. A) Depicts the perforation position of the maxillary suture of the rat: the two holes are about 12 mm apart. B) Sample location: 7 mm tissue of rat maxillary suture. C-D) Show a three-dimensional structure and planar structure of the stent. E) Shows a photo taken one week after the stent was placed in the rat. The stent was in the expected position and had maintained certain traction. F) Shows isolated specimens taken from the maxillary suture.

### 2.3 Production of traction devices

A 0.4 mm diameter titanium-nickel alloy wire was selected to make a W-shaped traction device that provides a traction of about 100 g. Utilizing the super-elastic characteristics of titanium-nickel memory alloy wire, it can generate a constant force value under constant deformation when placed in the rat.

### 2.4 Quantitative real-time PCR (QPCR)

First-strand cDNA synthesis was performed using an FastKing RT Kit (Tiangen, Dalian, China) and a high-capacity cDNA synthesis kit (Applied Biosciences) according to the manufacturer’s instructions, SYBR Green quantitative real-time reverse transcription PCR analysis was performed on a sequence detector (Applied Biosystems, Foster City, CA). For quantitative real-time PCR, a SYBR green kit (Tiangen, Dalian, China) was used, and the DNA was analyzed employing the ABI QPCR machine, following the manufacturer’s instructions. The PCR program used is as follows: start at 95 °C for 10 s, followed by 35 thermal cycles (95 °C for 5 s, 60 °C for 31 s). After QPCR, GAPDH was used as an internal reference control, and the relative fold change of expression was calculated by the 2^−(ΔΔCt)^ method[10]. All the primer sequences are listed in Supplementary Table S1. BLAST (National Institutes of Health, Bethesda, MD) was used to verify that all the primer sequences were specific for the targeted gene.

### 2.5 RNA quantification and qualification

The sacroiliac maxilla was exposed after general anesthesia at 1 and 2 weeks after surgery, and 7 mm bone tissue around the maxillary suture was taken (FIGURE 2. F). The collected samples were stored in liquid nitrogen. RNA degradation and contamination were monitored on 1% agarose gels. RNA purity was checked using the NanoPhotometer^®^ spectrophotometer (IMPLEN, CA, USA), and RNA concentration was measured using Qubit^®^ RNA Assay Kit with a Qubit^®^2.0 Fluorometer (Life Technologies, CA, USA). RNA integrity was assessed using the RNA Nano 6000 Assay Kit of the Bioanalyzer 2100 system (Agilent Technologies, CA, USA).

### 2.6 Library preparation for Transcriptome sequencing

A total amount of 3 µg RNA per sample was used as input material for sample preparation. Sequencing libraries were generated using NEBNext^®^ UltraTM RNA Library Prep Kit for Illumina® (NEB, USA) following manufacturer’s recommendations, and index codes were added to attribute sequences to each sample. Briefly, mRNA was purified from total RNA using poly-T oligo-attached magnetic beads. Fragmentation was carried out using divalent cations at an elevated temperature in an NEBNext First Strand Synthesis Reaction Buffer(5X). First strand cDNA was synthesized using a random hexamer primer and M-MuLV Reverse Transcriptase (RNase H^−^). Second strand cDNA synthesis was subsequently performed using DNA Polymerase I and Rnase H. The remaining overhangs were converted into blunt ends via exonuclease/polymerase activities. After adenylation of 3’ ends of the DNA fragments, NEBNext adaptors with hairpin loop structure were ligated to prepare for hybridization. To select cDNA fragments of preferentially 250∼300 bp in length, the library fragments were purified employing the AMPure XP system (Beckman Coulter, Beverly, USA). Next, 3 µl USER Enzyme (NEB, USA) was added to size-selected, adaptor-ligated cDNA, and the mixture was at 37 °C for 15 min, followed by 5 min at 95 °C. Subsequently, PCR was performed using Phusion High-Fidelity DNA polymerase, Universal PCR primers and Index (X) Primer. Finally, the PCR products were purified (AMPure XP system) and library quality was assessed on the Agilent Bioanalyzer 2100 system.

### 2.7 Clustering and sequencing (Novogene Experimental Department)

The clustering of the index-coded samples was performed on a cBot Cluster Generation System using TruSeq PE Cluster Kit v3-cBot-HS (Illumina) according to the manufacturer’s instructions. After cluster generation, the library preparations were sequenced on an Illumina Hiseq platform and 150 bp paired-end reads were generated.

### 2.8 Quality control

Among the 24 samples submitted, expz1w, expz2w, conz1w, and conz2w represent the 1-week traction group, the 2-week traction group, the 1-week control group, and the 2-week control group, respectively. In preliminary analysis of the samples, we found that reads of the expz1w_3 and expz2w_2 samples had poor correlation, and the distributions of the principal component analysis of the conz2w_4, con2w_6, and conz1w_4 reads were significantly different. After excluding these parts of the reads, the remaining data were retained (expz1w: 1/2/4/5/6, conz1w: 1/2/3/5/6, expz2w: 1/3/4/5/6, and conz2w: 1/2/3/5) for final data analysis.

### 2.9 RNA sequence analysis

Raw data were clipped by removing reads containing adapters, and those containing ploy-N and low-quality reads. All the downstream analyses were based on clean data with good quality. Sequences were aligned to the reference genome using Hisat2 v2.0.5[11]. FeatureCounts v1.5.0-p3 was used to count the reads numbers mapped to each gene[12], and the FPKM of each gene was calculated based on the length of the gene and read counts mapped to this gene.

### 2.11 Differential expression analysis

Differential expression analysis of two conditions/groups (two biological replicates per condition) was performed using the R package DESeq2 (1.16.1). Genes with an adjusted P-value <0.05 found by DESeq2 were designated as differentially expressed[13, 14]. Gene set enrichment analysis of differentially expressed genes was performed by the cluster Profiler R package[15] with Gene Ontology terms[16] and KEGG (http://www.genome.jp/kegg/).

### 2.13 Other statistical analysis

Means and standard deviation (SD) of the QPCR data were calculated. The Student-t test was used to determine if and to what extent there is a statistical difference between groups; p-value< 0.05 was considered statistically significant.

## 3. RESULTS

### 3.1 Differential expression analyses

Differential expression analyses were performed to evaluate the gene expression profiles of the zygomatic maxillary suture cells with/without traction. A total of 2754 genes were differentially expressed at the first week (expz1w vs. conz1w). Among them, 1777 genes were up-regulated and 977 genes were down-regulated (Figure 3A). A total of 3367 differentially expressed genes (DEGs) were identified by comparing the groups at two weeks. Among them, 1748 genes were up-regulated and 1691 genes were down-regulated (Figure 3B). Comparing the DEGs between the groups at 1 week and 2 weeks, 637 (35.85%) of the 1777 genes that were up-regulated in expz1w were down-regulated in expz2w, 977 genes that were down-regulated in week 1 had 236 genes up-regulated by 2 weeks, and 1495 new up-regulated genes appeared in expz2w (Figure 3C).

**FIGURE 3.**
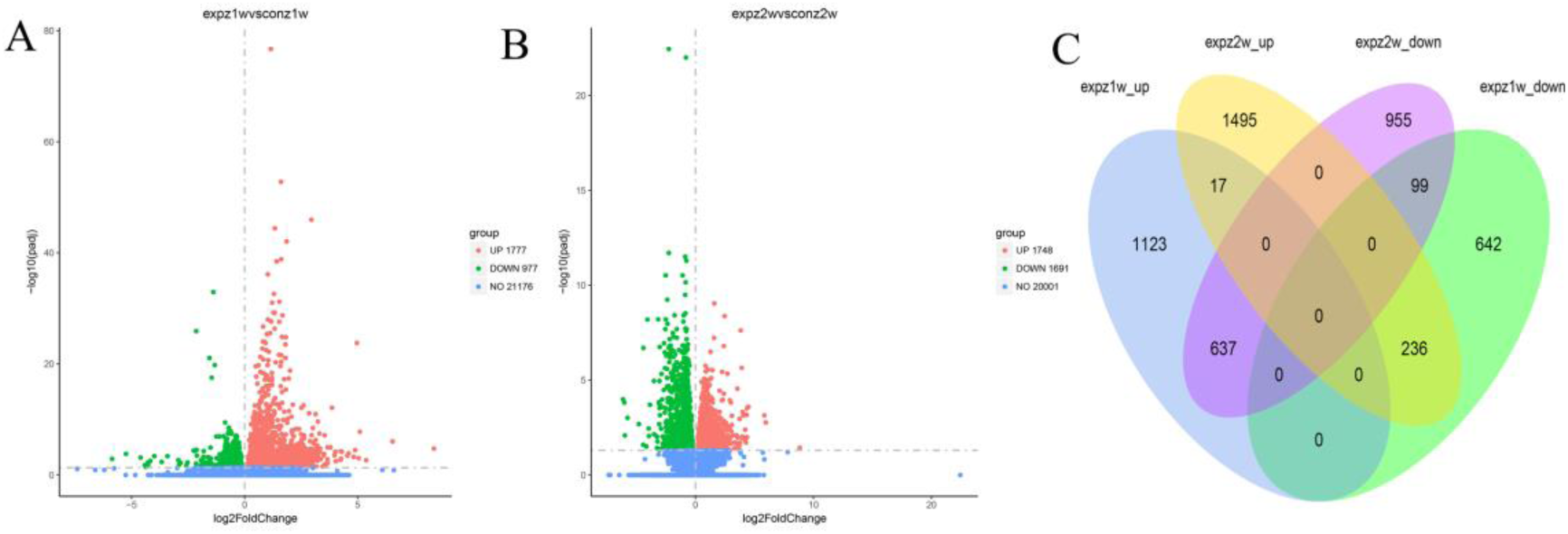
Volcano map and VENN chart of differentially expressed genes (DEGs). A) Volcano map of DEGs displaying a comparison between expz1w and conz1w. B) Volcano map of DEGs depicting a comparison between expz2w and conz2w. C) Venn diagram of DEGs.

### 3.2 Muscle development-related genes were significantly up-regulated at 1 week and down-regulated at 2 weeks

From the gene set enrichment analysis, we found that muscle-related genes had significant dynamic changes during traction. We found that 9 out of the top 10 biological process (BP) pathways in first week were related to promoting muscle system development. The top 10 cellular components (CC) in the first week were mainly related to cell collagen fiber production and cell matrix synthesis, while sarcomere, I Bands, Z discs, actin cytoskeleton, and basement membrane were closely related to skeletal muscle development. Actin binding, actin filament binding, extracellular matrix structural components, muscle structural components, extracellular matrix binding, and integrin binding in terms of molecular function (MF) pathways up-regulated in 1 week were also related to muscle development (Figure 4).

**FIGURE 4.**
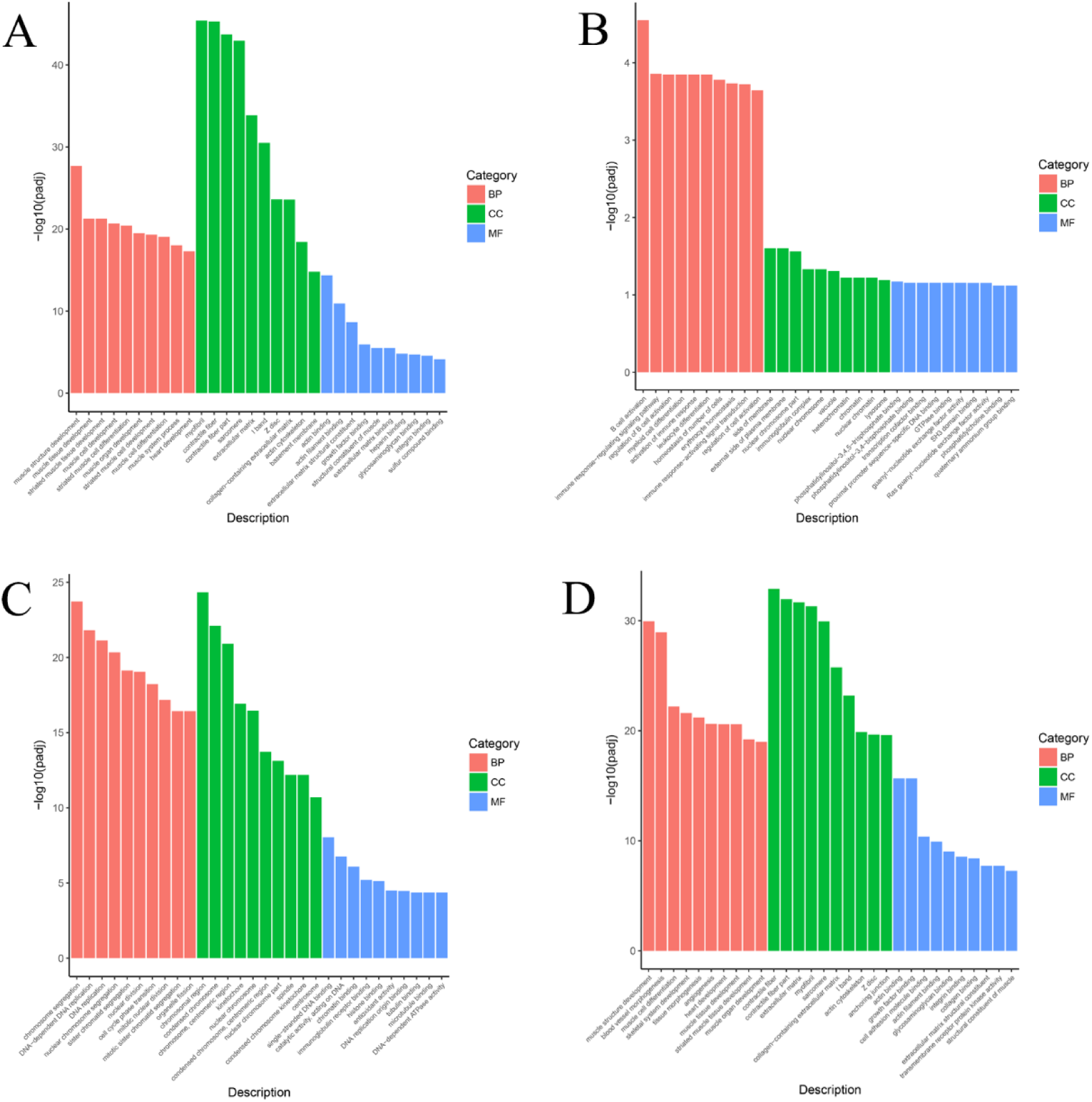
Gene ontology (GO) enrichment analysis of DEGs in the 1-week and 2-week groups. BP, Biological process; CC, Cellular components; MF, Molecular functions

In the top 10 BP pathways down-regulated in 2 weeks, six pathways were related to muscle development, including muscle cell differentiation, skeletal system development, muscle tissue development, striated muscle tissue development, and muscle organ development. Actin binding, actin filament binding, and muscle structural components of the MF pathway were all related to muscle development (Figure 4).

We sorted by the genes using log2 fold change to find the top 15 genes that were up-regulated and down-regulated at 2 weeks. Among them, *Xirp1, Abra, Myh3, Myl3, Ankrd1, Smpx*, and *MyBPC1* 6 genes (4<0.001%) were all related to muscle, and showed opposite trends at 1 and 2 weeks (P <0.05) (Table 2).

To study the relationship between the muscle-associated pathways that were up-regulated at 1 week and those that were down-regulated at 2 weeks, we separately calculated the Gene Ontology (GO) enrichment pathways for two weeks and we found that among the 827 GO enrichment pathways that were up-regulated at 1 week, 71 (8.59%) were related to muscle development. In addition, among the 1102 GO enrichment pathways down-regulated in 2 weeks, 101 (9.17%) were related to muscle development. From these data it was observed that muscle-related pathways accounted for an important proportion of both the 1-week and 2-week groups (Table 3). Further analysis revealed that 63 (62.38%) of 101 muscle-related pathways down-regulated at 2 weeks were the up-regulated muscle pathways at week 1 (Table 3). This indicates that muscle development-related genes were significantly up-regulated in the early stage of traction osteogenesis and were down-regulated at 2 weeks.

**TABLE 1.**
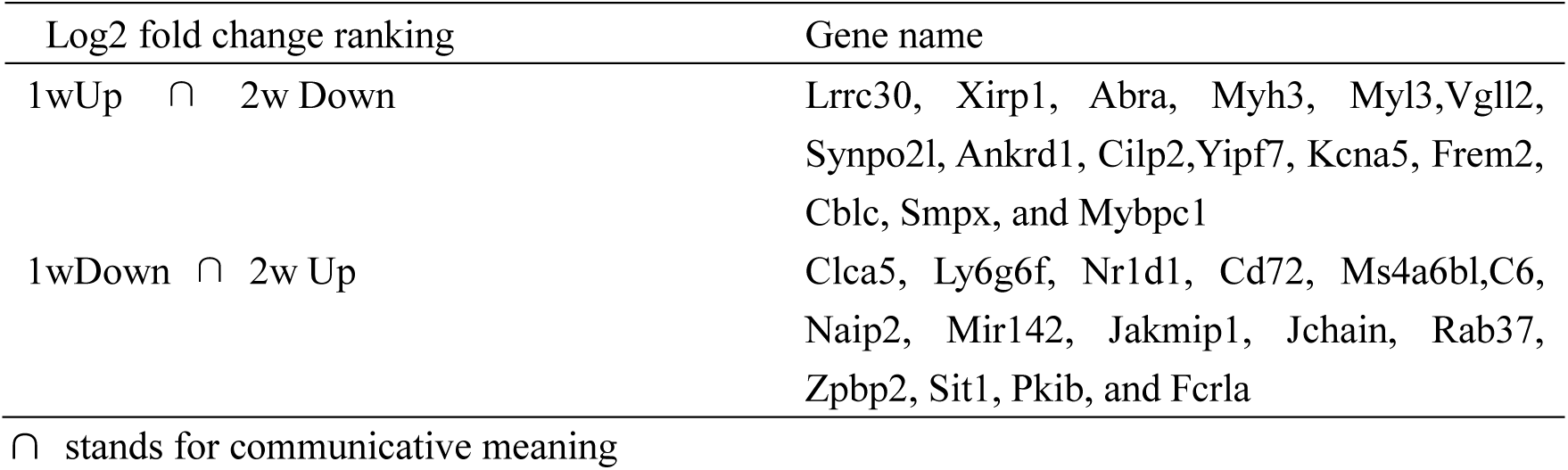
log2 fold change of 15 top-ranked genes.

**TABLE 3.** Muscle-related pathways in enriched Gene Ontology (GO) terms

|  | 1w Up | 1w Down | 2w Up | 2w Down |
| --- | --- | --- | --- | --- |
| BP | 68/659 | 0/114 | 0/589 | 98/949 |
| CC | 1/112 | 0/6 | 0/86 | 1/92 |
| MF | 2/56 | 0/0 | 0/50 | 2/61 |
| All | 71/827 | 0/120 | 0/725 | 101/1102 |

GO function enrichment analysis of the DEGs was further performed. The distribution of BP, CC, and MF pathways was in accordance with p-value, in the DEGS of the experimental as well as control groups. GO enrichment analysis was: A) increased at one week, B) reduced at one week, C) increased at two weeks, and D) reduced at two weeks.

### 3.3 The immune-related gene pathway was significantly down-regulated at 1 week, and the B-cell-related pathway was up-regulated at 2 weeks

*Ly6g6f, Cd72, C6, Naip2, Jchain, Fcrla*, and other genes among the 15 top-ranked genes that were down-regulated in the first week but up-regulated in the second week were all related to immunity (Table 2).The down-regulated GO terms at 1 week were B cell activation, immune response regulatory signaling pathway, B cell activation, bone marrow cell differentiation, activated immune response, white blood cell differentiation, homeostasis of cell number, and immune response. The immunoglobulin complexes in the CC pathway are immunologically related (Figure 4).

Further analysis of the immune-related pathways down-regulated at 1 week in the GO terms demonstrated that immune-related terms had an important role, of which 22 (18.33%) and immune-related pathways were found in 120 down-regulated pathways, while those up-regulated in 2 weeks. Among the 725 GO enrichment terms, 57 (7.86%) pathways were related to immunity. Immune-related pathways are of great significance in distraction osteogenesis (Table 4).

**TABLE 4.** B cell and immune pathways in the GO enrichment pathway.

|  | 1w Up | 1w Down | 2w Up | 2w Down |
| --- | --- | --- | --- | --- |
| BP | 0/659 | 22/114 | 57/589 | 0/949 |
| CC | 0/112 | 0/6 | 0/86 | 0/92 |
| MF | 0/56 | 0/0 | 0/50 | 0/61 |
| All | 0/827 | 22/120 | 57/725 | 0/1102 |

Further studies on the relevant immune pathways showed that B cells have an important role in immunity. Among the 22 immune pathways that were down-regulated in 1 week, 12 were related to B cells, while in the 57 immune pathways that were up-regulated in 2 weeks also 12 were related to B cells, of which 9 pathways were coincident (Table 5).

**TABLE 5.** B cell related pathways in 1w Down and 2w Up in Go enrichment analysis.

| GOID | Description | 1wDown |  | 2wUP |  |
| --- | --- | --- | --- | --- | --- |
|  |  | GeneRatio | padj | GeneRatio | padj |
| GO:0042113 | B cell activation | 36/757 | 2.81E-05 | 60/1403 | 1.75E-10 |
| GO:0050864 | regulation of B cell activation | 26/757 | <0.001 | 43/1403 | 1.71E-08 |
| GO:0050853 | B cell receptor signaling pathway | 19/757 | <0.001 | 33/1403 | 7.71E-08 |
| GO:0030183 | B cell differentiation | 14/757 | <0.001 | 15/1403 | 0.04 |
| GO:0050855 | regulation of B cell receptor signaling pathway | 5/757 | 0.01 | 5/1403 | 0.03 |
| GO:0042100 | B cell proliferation | 11/757 | 0.01 | 16/1403 | <0.001 |
| GO:0050871 | positive regulation of B cell activation | 17/757 | 0.02 | 35/1403 | 8.97E-07 |
| GO:0030888 | regulation of B cell proliferation | 9/757 | 0.02 | 12/1403 | <0.001 |
| GO:0019724 | B cell mediated immunity | 17/757 | 0.04 | 40/1403 | 1.71E-08 |

### 3.4 Cell growth- and development-related pathways show significant upregulation at 2 weeks

All the first 10 BP pathways that were up-regulated by GO enrichment analysis at 2 weeks were related to cell proliferation and replication, including chromosome isolation, DNA-dependent DNA replication, DNA replication, nuclear chromosome isolation, sister chromatid isolation, nuclear division, and cell cycle phase transitions, mitotic mitosis, mitotic sister chromatid separation, organelle fission. The CC pathway was related to chromosome region, chromosome enrichment, chromosome centromere region, spheroids, nuclear chromosome, concentrated chromosome centromere region, nuclear chromosome part, main axis, concentrated chromosome kinetics, and centrosome. The MF pathway was related to single-stranded DNA binding, catalytic activity on DNA, chromatin binding, immunoglobulin receptor binding, histone binding, antioxidant activity, DNA origin of replication binding, tubulin binding, microtubule binding, and DNA dependence ATPase activity (Figure 4), most of which were related to cell growth and development.

### 3.5 Up-regulation in several important pathways in traction osteogenesis by KEGG pathway analysis

The twenty top-ranking differential genes were analyzed by KEGG. The pathways that were up-regulated in 1 week included ECM-receptor interactions, focal adhesions, hippocampal signaling pathways-multiple species, PI3K-Akt signaling pathways, etc. (Fig 7), they are all related to suture osteogenesis. The signal pathway shows an upward trend. The B cell receptor signaling pathway in the down-regulated pathway was consistent with Go enrichment analysis. Among the top 20 pathways that were significantly up-regulated at 2 weeks were related to cell cycle, DNA replication, spliceosome, homologous recombination, mismatch repair, p53 signaling pathway, and cell senescence are all clearly related to cell growth and development. Adhesive plaques, ECM-receptor interactions, hippocampus signaling pathway, PI3K-Akt signaling pathway, hippocampus signaling pathway – a variety of species were significantly down-regulated at 2 weeks and were related to suture osteogenesis in the first week. Pathways, such as MAPK signaling pathways, and Wnt signaling pathways are newly emerging down-regulated cellular pathways (Figure 5).

**FIGURE 5.**
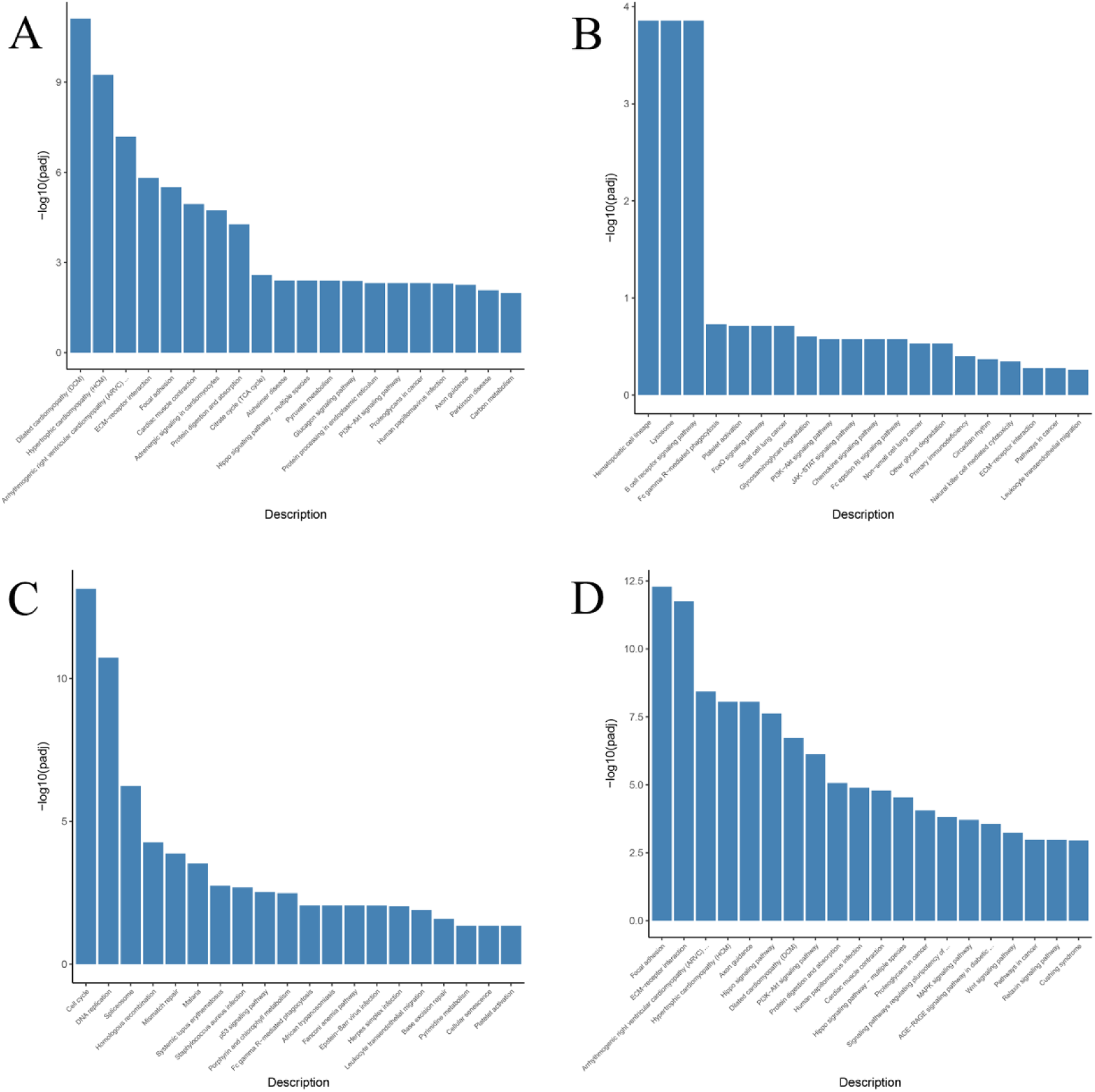
KEGG enrichment analysis of differential genes in the 1-week and 2-week groups. The differentially expressed genes were further subjected to KEGG function enrichment analysis, and then ranked according to padj. Pathways by KEGG enrichment analysis: A) increased at 1 week, B) decreased at 1 week, C) increased at 2 weeks. D) decreased at 2 weeks.

## 4 DISCUSSION

There is a close relationship between bones and muscles. Reports suggest that astronauts as well as patients who have been bedridden for a long time are prone to osteoporosis and muscle atrophy. Mechanical movement loads closely link bones and muscles. It is generally believed that muscles can promote bone healing due to its high vascularity, thus contributing to local blood supply. It is observed that ischemia can delay healing of fractures in mice. Besides, blood flow in the limbs is directly proportional to muscle mass; in addition, muscles provide a vascular supply to bones. However, fascial skin tissue with more blood vessels is not as effective as muscles in fracture healing [17-19]. Studies have reviewed the potential role of muscle in bone repair and have suggested that muscle-derived osteo-inductive cell populations may be directly related to bone formation and healing[20]. In our study, we also found that the genes that were up-regulated in the first week of TSDO surgery were mainly muscle-related genes. This indicates that muscle-related genes were mainly present in the early stage of traction and that muscles may be an initiating factor in the process of traction osteogenesis. Next, the mechanisms of interaction of muscles with bones were investigated. Some studies have found that muscles can secrete growth factors, cytokines and metallopeptidases. These proteins may be factors that interact with muscles and bones[21]. Researchers have identified some proteins secreted by muscles related to extracellular matrix remodeling, cell proliferation, migration, and signaling, and found that these molecules interact to play a functional role in myogenic differentiation[22]. Such factors include, insulin-like growth factor-1 (IGF-1), basic fibroblast growth factor (FGF-2), IL-6, IL-15, myostatin, osteoglycoprotein (OGN), FAM5C, Tmem119 and osteoactivin. Prostaglandin E2 (PGE2) and Wnt3a secreted by osteocytes, osteocalcin (OCN), and IGF-1 secreted by osteocytes may affect skeletal muscle cells. In addition, chondrocytes secrete Dickkopf-1 (DKR-1) and India hedgehog(Ihh), while adipocytes produce leptin, adiponectin, and IL-6, which can also regulate bone and muscle metabolism[23].

The concept of osteoimmunology recently been proposed by investigators suggests that immune cells play an important role in controlling the homeostasis of bones[24]. In our study, we found that the first 10 BP pathways down-regulated by GO enrichment analysis neutralized more immune-related pathways in these pathways, including B cell activation, immune response regulatory signaling pathways, B cell activation regulation, bone marrow cell differentiation, and activation. Immune response, leukocyte differentiation, and immune response activation signal transduction are all related to immunity. Previous studies have found that B cells play an important role in bone immunity. Early B cells are produced by the bone marrow and share the same cytokines, chemokines, receptors and downstream signaling pathways as early osteocalcin (OC) or osteoblast (OB). The B cell family (including precursor B cells, immature B cells, and stromal cells) in the bone marrow of mice produces 64% of OPG in the bone marrow, a finding that has revolutionized the study of early osteoimmunology. In bone immunology, cells of the immune system regulate bone immunity through complex interactions with RANK / RANKL / OPG axis. Studies have found that OPG produced by B cells is beneficial to maintain a balance between RANKL / RANK / OPG, thus maintaining the balance of bone metabolism[25]. Changes in B cells and T cells in human immunodeficiency virus (HIV)-infected persons can affect the osteoclast receptor activators of OPG and RANKL cytokines secreted by OBs, thereby promoting osteoclastogenesis and bone resorption and leading to bone loss in patients[26]. Studies have found that B cells can support OC differentiation *in vitro* in a RANKL-dependent manner and that the number of OC cultured from patients with rheumatoid arthritis is greater than that in healthy controls[27]. In addition, clinical studies have found that in menopausal women with osteoporosis, the numbers of CD19 + B cells and memory B cell subpopulations are significantly reduced compared to those in healthy controls[28]. Some studies have analyzed the genomic expression levels in menopausal women with osteoporosis and ovariectomized mouse models through microarray technology. Genes in these women and the mouse models exhibited similar changes, including many genes related to B cell function and development. These results suggest that some of the pathogenesis of bone loss and osteoporosis diseases may be related to B cell abnormalities[29]. In our study too, we focused on the role of multiple B cell-related pathways in distraction osteogenesis, but the specific mechanisms remain to be further explored.

Macrophages are the primary regulatory cells for the immune response during osteogenesis and angiogenesis. Studies have shown that macrophages can reduce the expression of inflammatory factors, such as TNF-α and IL-6 and are related to the up-regulation of TGF-β1 related to bone repair[30, 31]. Two types of immune cytokines work together to regulate the balance of bone metabolism. Among them, protective cytokines, including IL-2, IL4, IL-10, IL-12, IL-13, IL-18, IFN, tend to positively balance bone metabolism, can block RANKL signal, directly or indirectly inhibit bone resorption, and promote bone formation; whereas other inflammatory cytokines, such as IL-6, TNF, IL-1, IL-3, IL-7, IL-11, IL-15 and IL-17 tend to up-regulate RANKL expression, promote negative bone metabolism balance, enhance osteoclast activity, and promote bone resorption [32]. There are two subtypes of tumor necrosis factor, α and β. TNF-α causes bone resorption and inhibits arthritis and bone formation, and TNFβ can stimulate bone marrow stromal cells to express RANKL, activate p38MAPK signaling pathway, increase M-CSF receptor; C-fms gene expression further promotes OC generation.

Mesenchymal stem cells (MSCs) are a type of cells that play an important role in the process of osteogenesis in suture traction. Some investigators have demonstrated that MSCs present in long bones and skulls have clonal pluripotency and self-renewal ability[33]. Recently, some researchers have demonstrated that Gli1 + cells are the main cells present in the MSC population of the cranial sutures of mice. These cells have the potential to generate craniofacial bone and can be activated again during craniofacial bone repair. The loss of Gli1 + cells can cause a loss of cranial sutures. These cells are indispensable stem cell populations in the growth and development of the skull and in repair of injuries[7]. In the process of traction osteogenesis, different signal pathways are activated, and their interactions form a network, leading to MSC proliferation and differentiation and ultimately jointly regulating the process of traction osteogenesis. In our research, we found that the cell cycles related to the regulation of growth and development appeared to be significantly up-regulated during the two weeks of traction. Among them, pathways related to chromosome separation, DNA-dependent DNA replication, DNA replication, nuclear chromosome separation, sister chromatid separation, nuclear division, and cell cycle phase transition were found. In addition, pathways, such as mitotic mitosis, mitotic sister chromatid separation, and organelle fission, closely related to cell proliferation and differentiation were also up-regulated.

During traction osteogenesis, mechanical traction force is transformed into biomolecular signals, causing bone suture tissue cells to secrete bone morphogenetic proteins (BMPs), platelet-rich plasma (PRP), transforming growth factor β (TGF-β), and other growth factors Under the control of the growth factor network, mesenchymal stem cells (MSCs) proliferate and differentiate into osteoblasts and chondrocytes, and form new bone through osteogenesis, such as intra-membrane osteogenesis and intra-chondral osteogenesis[5, 34]. This process involves the activation of a variety of cellular pathways. Previous studies have found that integrin signaling pathway, mTOR signaling pathway, Wnt / β-catenin signaling pathway, Hippo pathway, and others, play important roles in traction osteogenesis. Some studies have found that Smad pathway, mitogen-activated protein kinase (MAPKs) pathway, etc., are also activated to varying degrees in osteotomy osteogenesis[35-38]. In our study, we found that ECM-receptor interaction, focal adhesion, Hippo signaling pathway-multiple species, PI3K-Akt signaling pathway, Wnt signaling pathway, Hippo signaling pathway, and other pathways show obvious trends of up-regulation in traction osteogenesis, thus indicating that these pathways may indeed be important in distraction osteogenesis.

## ACKNOWLEDGMENTS

The project was supported by the National Natural Science Foundation of China (No. 81571925)

## AUTHOR CONTRIBUTIONS

All authors took part in writing, reviewing, and editing the manuscript, ZN Zhao, BC Wang, HZ Tong and HY Xue designed the experiments; BC Wang, HZ Tong, YD Sun, PY Zhang performed the experiments and prepared figures. All the authors reviewed the manuscript and approved it for publication.

## Supplementary material

### 1. Sample quality control

**FIGURE S1.**
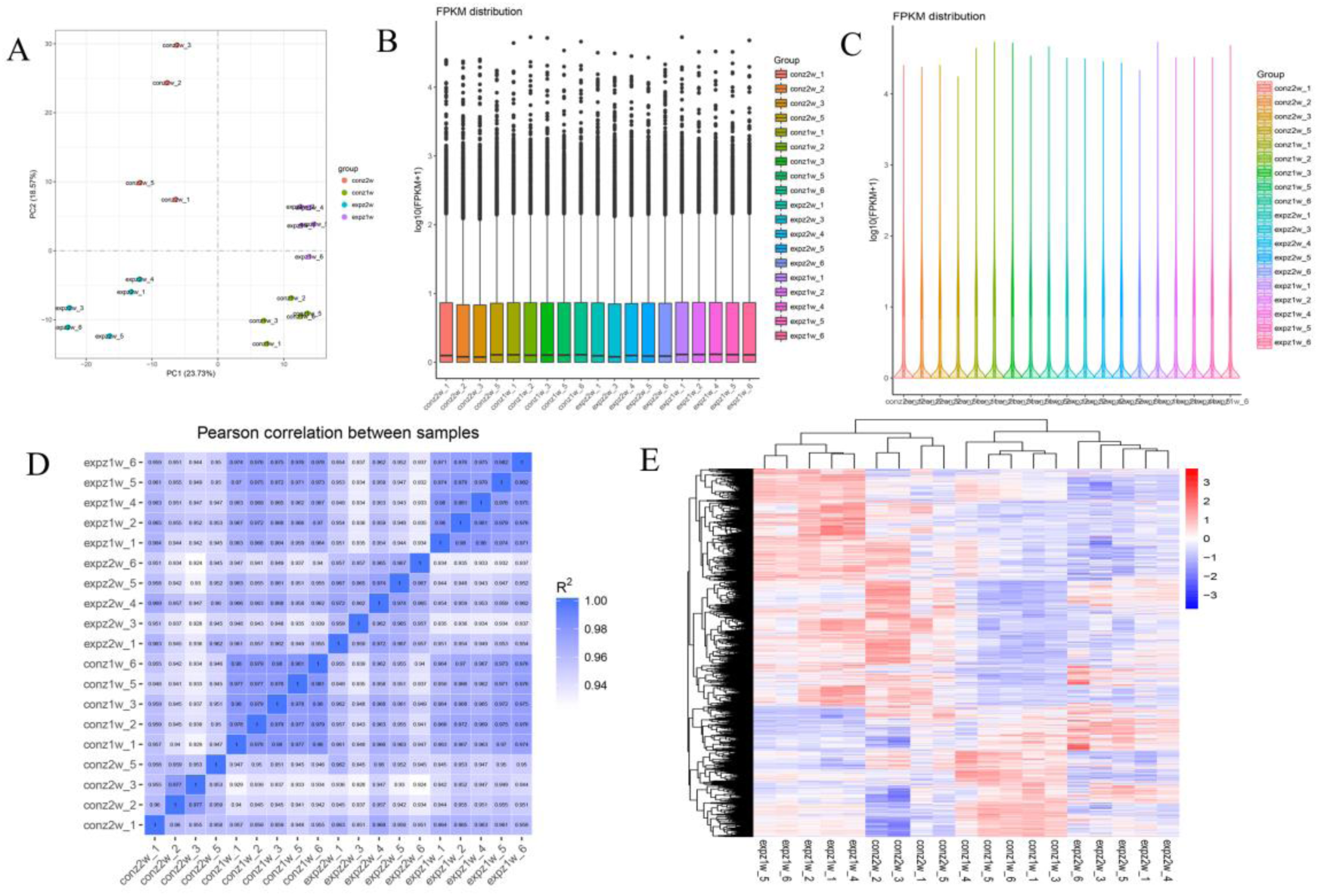
Data quality control. A) shows all of the samples on the first (x-axis) and second (y-axis) principal component. B-C) show the library size of the samples. D) The square (R2) of the Pearson correlation coefficient in this experiment is>0.94. E) Clusters of the differential gene sets related to cluster genes with similar expression patterns in the whole transcriptome.

### 2. QPCR results

The selected differential genes (Fat4, Yap1, Sema3C, Dchs1, and Gli1) were validated by QPCR. The verified results were consistent with the whole transcriptome analyses.

**TABLE S1.** Genes up-regulated at one week after TSDO.

| gene name | Expz FPKM | Conz FPKM | log2 fold change | Adjusted p-value |
| --- | --- | --- | --- | --- |
| Fat4 | 1292.15 | 579.07 | 1.15 | 1.99E-77 |
| Yap1 | 1060.54 | 779.97 | 0.44 | 4.91E-05 |
| Sema3c | 454.09 | 125.58 | 1.85 | 8.89E-43 |
| Dchs1 | 2650.61 | 2012.84 | 0.39 | $< 0.001$ |
| Gli1 | 641.90 | 442.16 | 0.53 | $< 0.001$ |

**FIGURE S2.**
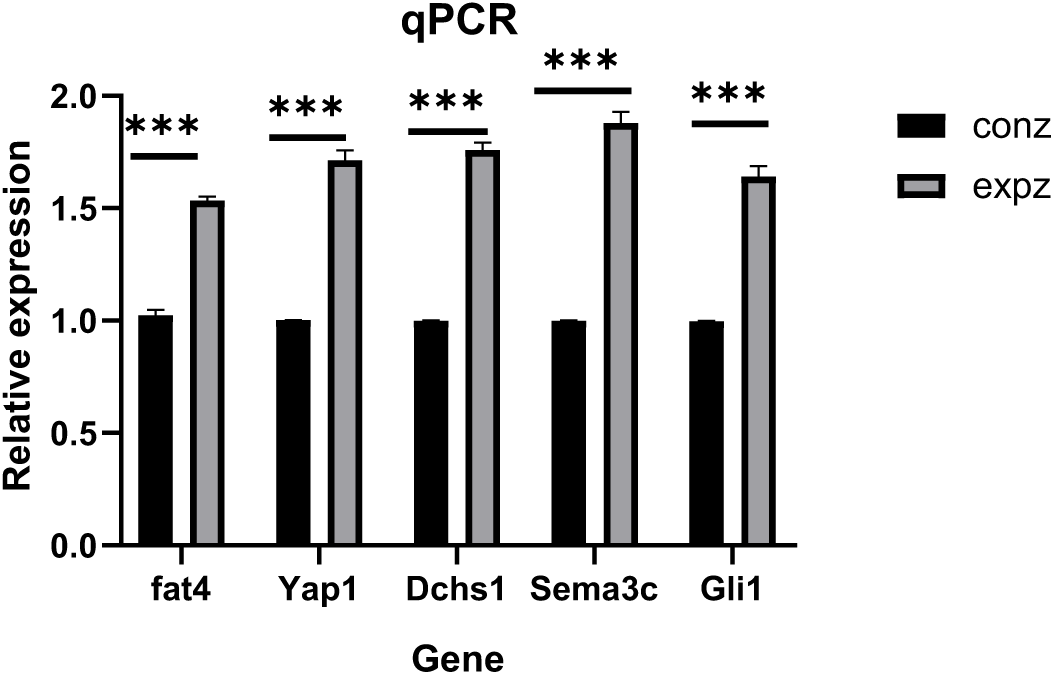
QPCR results of up-regulation of some genes in one week, *** significant difference

**TABLE S2.**
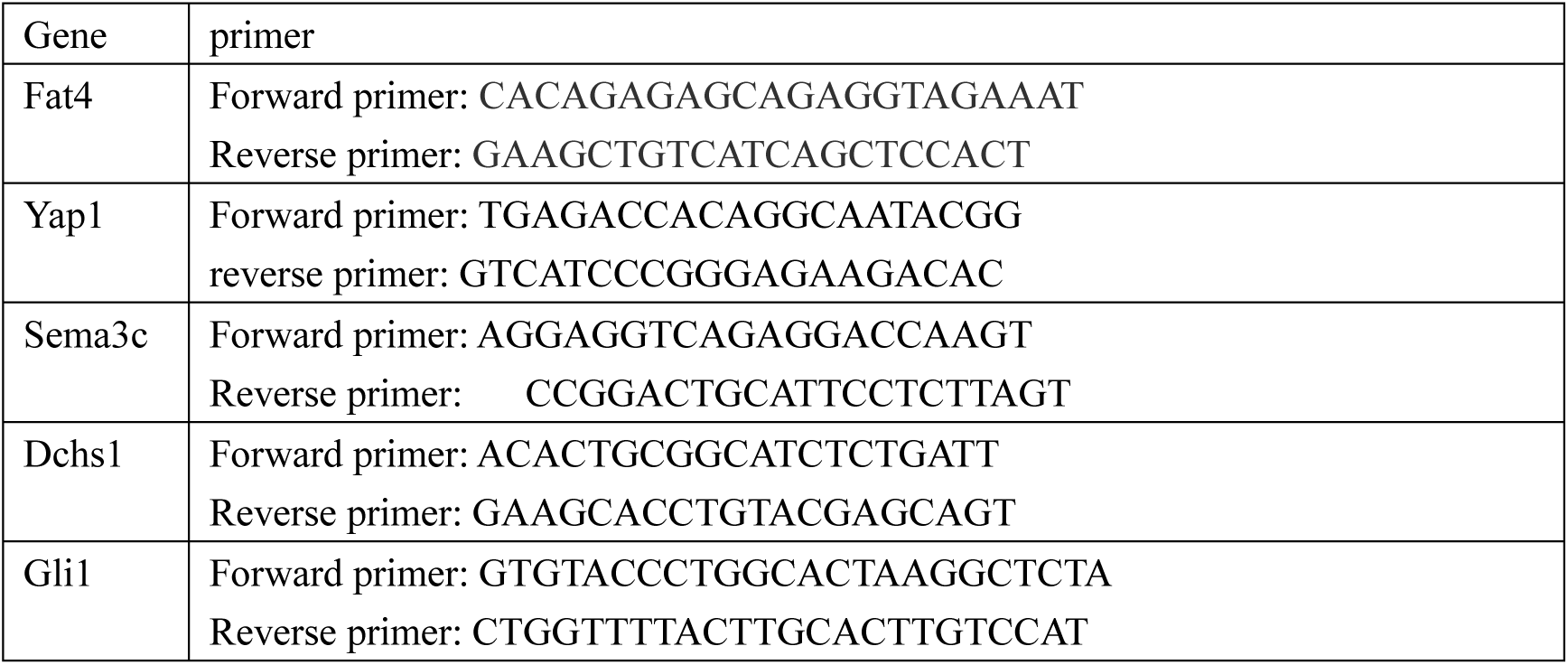
PCR primers.

